# A Fast Data-Driven Method for Genotype Imputation, Phasing, and Local Ancestry Inference: MendelImpute.jl

**DOI:** 10.1101/2020.10.24.353755

**Authors:** Benjamin B. Chu, Eric M. Sobel, Rory Wasiolek, Janet S. Sinsheimer, Hua Zhou, Kenneth Lange

## Abstract

Current methods for genotype imputation and phasing exploit the sheer volume of data in haplotype reference panels and rely on hidden Markov models. Existing programs all have essentially the same imputation accuracy, are computationally intensive, and generally require pre-phasing the typed markers. We propose a novel data-mining method for genotype imputation and phasing that substitutes highly efficient linear algebra routines for hidden Markov model calculations. This strategy, embodied in our Julia program `MendelImpute.jl`, avoids explicit assumptions about recombination and population structure while delivering similar prediction accuracy, better memory usage, and an order of magnitude or better run-times compared to the fastest competing method. `MendelImpute` operates on both dosage data and unphased genotype data and simultaneously imputes missing genotypes and phase at both the typed and untyped SNPs. Finally, `MendelImpute` naturally extends to global and local ancestry estimation and lends itself to new strategies for data compression and hence faster data transport and sharing.

## 2 Introduction

Haplotyping (phasing) is the process of inferring unobserved haplotypes from observed genotypes. It is possible to deduce phase from the observed genotypes of surrounding pedigree members [25], but pedigree data are no longer considered competitive with linkage disequilibrium data. Current methods for phasing and genotype imputation exploit public reference panels such as those curated by the Haplotype Reference Consortium (HRC) [23] and the NHLBI TOPMed Program [27]. The sizes of these reference panels keep expanding: from 1000 samples in 2012 [2], to about 30,000 in 2016 [23], and to over 90,000 in 2019 [27]. Genome-wide association studies (GWAS), the primary consumers of imputation, exhibit similar trends in increasing sample sizes and denser SNP typing [26]. Despite these technological improvements, phasing and imputation methods are still largely based on hidden Markov models (HMM). Through decades of successive improvements, HMM software is now more than 10,000 times faster than the original software [10], but the core HMM principles remain relatively unchanged. This paper explores an attractive data-driven alternative for imputation and phasing that is faster and simpler than HMM methods.

Hidden Markov models (HMMs) capture the linkage disequilibrium in haplotype reference panels based on the probabilistic model of Li and Stephens [21]. The latest HMM software programs include `Minimac` 4 [11], `Beagle` 5 [7], and `Impute` 5 [24]. These HMMs programs all have essentially the same imputation accuracy [7], are computationally intensive, and generally require pre-phased genotypes. The biggest computational bottleneck facing these programs is the size of the HMM state space. An initial pre-phasing (imputation) step fills in missing phases and genotypes at the typed markers in a study. The easier second step constructs haplotypes on the entire set of SNPs in the reference panel from the pre-phased data [14]. This separation of tasks forces users to chain together different computer programs, reduces imputation accuracy [10], and tends to inflate overall run-times even when the individual components are well optimized.

Purely data-driven techniques are potential competitors to HMMs in genotype imputation and haplotyping. Big data techniques substitute massive amounts of training data for detailed models in prediction. This substitution can reduce computation times and, if the data are incompatible with the assumptions underlying the HMM, improve accuracy. Haplotyping HMMs, despite their appeal and empirically satisfying error rates, make simplifying assumptions about recombination hot spots and linkage patterns. We have previously demonstrated the virtues of big data methods in genotype imputation with haplotyping [8] and without haplotyping [9]. `SparRec` [15] refines the later method by adding additional information on matrix co-clustering. These two matrix completion methods efficiently impute missing entries via low-rank approximations. Unfortunately, they also rely on computationally intensive cross validation to find the optimal rank of the approximating matrices. On the upside, matrix completion circumvents pre-phasing, exploits reference panels, and readily imputes dosage data, where genotype entries span the entire interval [0, 2].

Despite these advantages, data-driven methods have not been widely accepted as alternatives to HMM methods. Although it is possible in principle, our previous program [9] did not build a pipeline to handle large reference panels. Here we propose a novel data-driven method to fill this gap. Our software `MendelImpute` (a) avoids the pre-phasing step, (b) exploits known haplotype reference panels, (c) supports dosage data, (d) runs extremely fast, (e) makes a relatively small demand on memory, and (f) naturally extends to local and global ancestry inference. Its imputation error rate is slightly higher than the best HMM software but still within a desirable range. `MendelImpute` is open source and forms a part of the `OpenMendel` platform [28], which is in the modern Julia programming language [4]. We demonstrate that `MendelImpute` is capable of dealing with HRC data even on a standard laptop. In coordination with our packages `VCFTools.jl` (handling VCF files) and `SnpArrays.jl` (handling PLINK files), `OpenMendel` powers a streamlined pipeline for end-to-end data analysis.

For each chromosome of a study subject, `MendelImpute` reconstructs two extended haplotypes **E**_1_ and **E**_2_ that cover the entire chromosome. Both **E**_1_ and **E**_2_ are mosaics of reference haplotypes with a few break points where a switch occurs from one reference haplotype to another. The break points presumably represent contemporary or ancient recombination events. `MendelImpute` finds these reference segments and their break points. From **E**_1_ and **E**_2_ it is trivial to impute missing genotypes, both typed and untyped. The extended haplotypes can be painted with colors indicating the region on the globe from which each reference segment was drawn. The number of SNPs assigned to each color immediately determine ethnic proportions and plays into admixture mapping. The extended segments also serve as a convenient device for data compression. One simply stores the break points and the index of the reference haplotype assigned to each segment. Finally, **E**_1_ and **E**_2_ can be nominated as maternal or paternal whenever either parent of a sample subject is also genotyped.

## 3 Materials and Methods

Our overall imputation strategy operates on an input matrix **X** whose columns are sample genotypes at the typed markers. The entries of **X** represent alternative allele counts *x_i j_* ∈ [0, 2] ∪ {missing}. The reference haplotypes are stored in a matrix **H** whose columns are haplotype vectors with entries *h_i j_* ∈ {0, 1}, representing reference and alternative alleles, respectively. Given these data, the idea is to partition each sample’s genotype vector into small adjacent genomic windows. In each window, many reference haplotypes collapse to the same unique haplotype at the typed SNPs. We find the two unique haplotypes whose vector sum best matches the sample genotype vector. Then we expand the unique haplotypes into two sets of matching full haplotypes and intersect these sets across adjacent windows. Linkage disequilibrium favors long stretches of single reference haplotypes punctuated by break points. Our strategy is summarized in Figure 1. A detailed commentary on the interacting tactics appears in subsequent sections.

**Figure 1:**
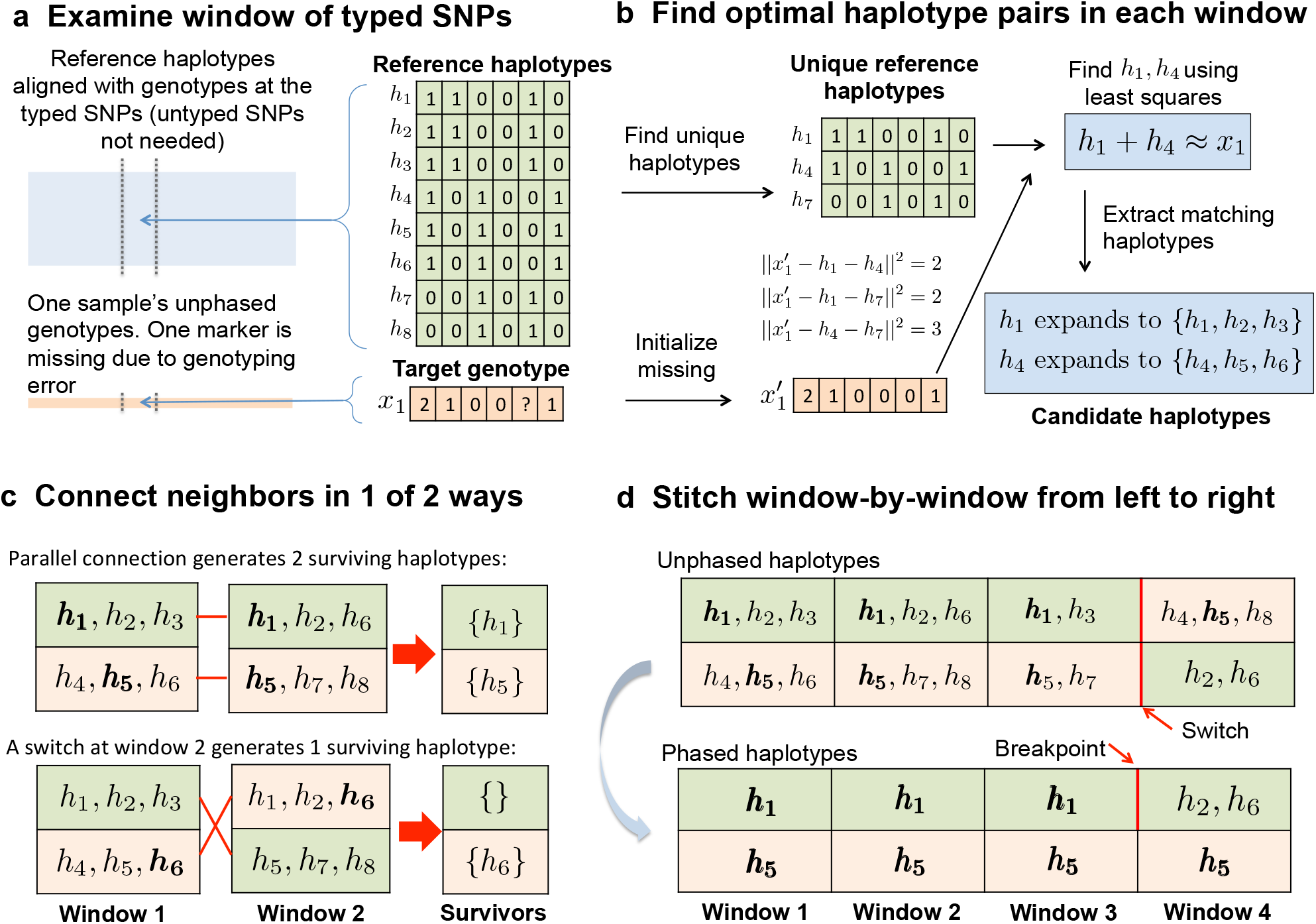
Overview of `MendelImpute`’s algorithm. (a) After alignment, imputation and phasing are carried out on short, non-overlapping windows of the typed SNPs. (b) Based on a least squares criterion, we find two unique haplotypes whose vector sum approximates the genotype vector on the current window. Once this is done, all reference haplotypes corresponding to these two unique haplotypes are assembled into two sets of candidate haplotypes. (c) We intersect candidate haplotype sets window by window, carrying along the surviving set and switching orientations if the result generates more surviving haplotypes. (d) After three windows the top extended chromosome possess no surviving haplotypes, but a switch to the second orientation in the current window allows **h_5_** to survive on the top chromosome. Eventually we must search for a break point separating **h_1_** from **h_2_** or **h_6_** between windows 3 and 4 (bottom panel).

### 3.1 Missing Data in Typed and Untyped SNPs

There are two kinds of missing data requiring imputation. A GWAS data set may sample on the order of 10^6^ SNPs across the genome. We call SNPs that are sampled at this stage typed SNPs. Raw data from a GWAS study may contain entries missing at random due to experimental errors, but the missing rate is usually low, at most a few percent, and existing programs [22] usually impute these in the pre-phasing step. When modern geneticists speak of imputation, they refer to imputing phased genotypes at the unsampled SNPs present in the reference panel. We call the unsampled markers untyped SNPs. The latest reference panels contain from 10^7^ to 10^8^ SNPs, so an imputation problem can have more than 90% missing data. We assume that the typed SNPs sufficiently cover the entire genome. From the mosaic of typed and untyped SNPs, one can exploit local linkage disequilibrium to infer for each person his/her phased genotypes at all SNPs, typed and untyped. As a first step one must situate the typed SNPs among the ordered SNPs in the reference panel (Figure 1A). The Julia command `indexin`() quickly finds the proper alignment.

### 3.2 Finding Optimal Haplotype Pairs via Least Squares

Suppose there are *d* unique haplotypes **h**_1_, …, **h**_*d*_ (entries 0 or 1) in a genomic window (Figure 1B), where the supplemental discusses how to efficiently compute them. Consider a genotype vector **x**with entries *x_i_* ∈ [0, 2] ∪ {missing}. The goal is to find the two unique haplotypes **h**_*i*_ and **h**_*j*_ such that **x**≈ **h**_*i*_ + **h**_*j*_. The best haplotype pair is selected by minimizing the least squares criterion

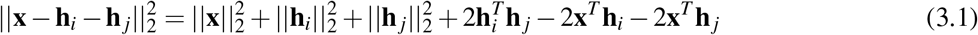

over all 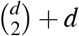 haplotype combinations. To fill in a missing value *x_i_*, we naively initialize it with the mean at each typed SNP. This action may lead to imputation errors if the typed SNPs exhibit a large proportion of missingness. We discuss a strategy to remedy this bias in the supplemental.

To minimize the criterion (3.1) efficiently, suppose the genotype vectors **x**_*i*_ constitute the columns of a genotype matrix **X**, and suppose the haplotype vectors **h**_*i*_ constitute the columns of a haplotype matrix **H**. Given these conventions we recover all inner products 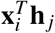 and 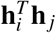 in equation (3.1) as entries of two matrix products; the two corresponding BLAS (Basic Linear Algebra Subroutines)[19] level-3 calls produce

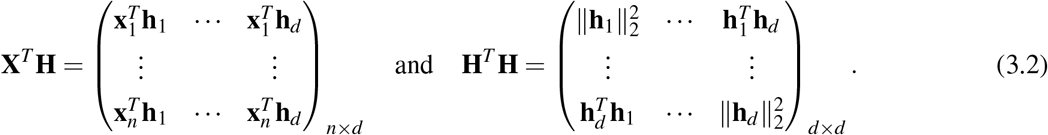

These allow one to quickly assemble a matrix **M** with entries 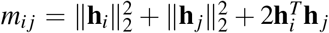 and for each sample **x**_*k*_ a matrix **N** with entries 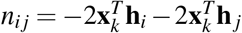. Therefore, to find the best haplotype pair (**h**_*i*_, **h**_*j*_) for the sample **x**_*k*_, we search for the minimum entry *m_i__j_* + *n_i__j_* of the *d* × *d* matrix **M** + **N** across all indices *i* ≥ *j*. Extremely unlikely ties are arbitrarily broken. Note that the constant term 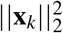 can be safely ignored in optimization. Data import and this minimum entry search are the computational bottlenecks of our software. Once such a haplotype pair is identified, all reference haplotype pairs identical to (**h**_*i*_, **h**_*j*_) in the current window give the same optimal *ℓ*_2_ error.

### 3.3 Phasing by Intersecting Haplotype Sets

As just described, each window *w* along a sample chromosome generates an optimal pair of unique haplotypes. These expand into two sets *S_w1_* and *S_w2_* of reference haplotypes (Figure 1B). In the first window we arbitrarily assign *S*_11_ to extended haplotype 1 and *S*_12_ to extended haplotype 2. From here on the goal is to reconstruct two extended composite haplotypes **E**_1_ and **E**_2_ that cover the entire chromosome. Let *w* index the current window. The two sets *S_w−1,1_* and *S_w−1,2_* are already phased. The new sets *S_w1_* and *S_w2_* are not, and their phases must be resolved and their entries pruned by intersection to achieve extended haplotype parsimony. The better orientation is one which generates more surviving haplotypes after intersection (Figure 1C). Here we count surviving haplotypes across both sets of an orientation. The better orientation and the corresponding survivor sets are propagated to subsequent windows. If either intersection is empty at window *w*, then a break is declared, the empty set is replaced by the entire haplotype set of window *w*, and a new reference segment commences (Figure 1D). Ties and double empties virtually never occur. Repeated intersection may fail to produce singleton haplotype sets, in which case we randomly designate a winner to use for breakpoint search.

For example, suppose *S*_11_ = {**h**_1_, **h**_2_, **h**_3_} and *S*_12_ = {**h**_4_, **h**_5_, **h**_6_} are the (arbitrarily) phased sets in window 1. Since window 2 is not yet phased, the two sets *S*_21_ = {**h**_1_, **h**_2_, **h**_6_} and *S*_22_ = {**h**_5_, **h**_7_, **h**_8_} can be assigned to extended haplotypes 1 and 2, respectively, or vice versa as depicted in Figure 1C. The first orientation is preferred since it generates two surviving haplotypes **h**_1_ and **h**_5_ bridging windows 1 and 2. Thus, {**h**_1_} and {**h**_5_} are assigned at window 2 with this orientation and propagated to window 3. In window 3 the contending pairs are {**h**_1_} ∩ {**h**_1_, **h**_3_} and {**h**_5_} ∩ {**h**_2_, **h**_5_} versus {**h**_1_} ∩ {**h**_2_, **h**_5_} and {**h**_5_} ∩ {**h**_1_, **h**_3_}. The former prevails, and {**h**_1_} and {**h**_5_} are assigned to window 3 and propagated to window 4. In window 4 the opposite orientation is preferred (Figure 1D). In this empty intersection case we set *S*_41_ = {**h**_2_, **h**_6_} and *S*_42_ = {**h**_5_} and continue the process. Later we return and resolve the breakpoint in extended haplotype 1 between windows 3 and 4.

### 3.4 Resolving Breakpoints

The unique haplotype pairs found for adjacent windows are sometimes inconsistent and yield empty intersections. In such situations, we search for a good break point. Figure 1D illustrates a single-breakpoint search. In this example, we slide the putative break point *b* across windows 3 and 4 in the top extended haplotype to minimize the least squares value determined by the observed genotype, **h**_5_ spanning both windows, and the breakpoint *b* between **h**_1_ and **h**_2_ ∪ **h**_6_. When there is a double mismatch, we must search for a pair (*b*_1_*, b*_2_) of breakpoints, one for each extended haplotype. The optimal pair can be determined by minimizing the least squares distances generated by all possible breakpoint pairs (*b*_1_*, b*_2_). Thus, double breakpoint searches scale as a quadratic. Fortunately, under the adaptive window width strategy described in Section 6.2.1, the number of typed SNPs in each window typically is on the order of 10^2^. In this range, quadratic search remains fairly efficient.

### 3.5 Imputation and Phasing of Untyped SNPs

Once haplotyping is complete, it is trivial to impute missing SNPs. Each missing SNP is located on the reference map, and its genotype is imputed as the sum of the alleles on the extended haplotypes **E**_1_ and **E**_2_. Observed genotypes are untouched unless the user prefers phased genotypes. In this case `MendelImpute` will override observed genotypes with phased haplotypes similar to `Minimac` 4. Unfortunately, `MendelImpute` cannot compute estimated dosages. As shown in Section 4.4 on alternative compression schemes, the extended haplotypes **E**_1_ and **E**_2_ can be output rather than imputed genotypes at the user’s discretion.

### 3.6 Compressed Reference Panels

Large reference files are typically stored as compressed VCF files. Since VCF files are plain text files, they are large and slow to read. Read times can be improved by computing and storing an additional tabix index file [20], but file size remains a problem. Consequently, every modern imputation program has developed its own specialized reference file format (for instance, the `m3vcf`, `imp5`, and `bref3` formats of `Minimac, Impute`, and `Beagle`, respectively) for improving read times and storage efficiency. We propose yet another compressed format for this purpose: the `jlso` format, and we compare it against other formats in Table 1. Details for generating the `jlso` format are discussed in the supplemental.

**Table 1:** Storage size required for various compressed reference haplotype formats. Here `vcf.gz` is the standard compressed VCF format, jlso is used by `MendelImpute`, `bref3` is used by `Beagle` 5.1, `m3vcf.gz` is used by `Minimac` 4, and imp5 is used by `Impute` 5. For all jlso files we chose the maximum number of unique haplotypes per window to be *d_max_* = 1000. Note we could not generate the `m3vcf.gz` file for the sim 1M panel because it required too much memory (RAM).

| Data Set | size (MB) for format: |  |  |  |  |
| --- | --- | --- | --- | --- | --- |
|  | vcf.gz | jlso | bref3 | m3vcf.gz | imp5 |
| sim 10K | 49 | 6 | 18 | 7 | 57 |
| sim 100K | 472 | 56 | 81 | 29 | 565 |
| sim 1M | 4660 | 776 | 415 | NA | 5650 |
| 1000G chr10 | 459 | 188 | 342 | 116 | 621 |
| 1000G chr20 | 200 | 89 | 154 | 52 | 279 |
| HRC chr 10 | 3346 | 1328 | 1156 | 529 | 3636 |
| HRC chr 20 | 1510 | 706 | 554 | 253 | 1610 |

### 3.7 Real and Simulated Data Experiments

For each data set, we exclude any typed SNPs with fewer than 5 copies of the minor allele and use only bi-allelic SNPs. Table 2 summarizes the real and simulated data used in our comparisons.

**Table 2:** Summary of real and simulated data sets used in our experiments. For the 1000G and HRC datasets, the SNPs also present on the Infinium Omni5-4 Kit constitute the typed SNPs. Missing % is the percentage of typed SNPs randomly masked to mimic random genotyping error. (* We used the top 50,000 most ancestry informative SNPs.)

| Data Set | Total SNPs | Typed SNPs | Samples | Ref Haplotypes | Missing % |
| --- | --- | --- | --- | --- | --- |
| Sim 10K | 62704 | 22879 | 1000 | 10000 | 0.5 |
| Sim 100K | 80029 | 23594 | 1000 | 100000 | 0.5 |
| Sim 1M | 97750 | 23092 | 1000 | 1000000 | 0.5 |
| 1000G Chr10 | 1511445 | 158569 | 100 | 4808 | 0.1 |
| 1000G Chr18 | 852602 | 50000* | 100 | 4808 | 0.1 |
| 1000G Chr20 | 669987 | 92652 | 100 | 4808 | 0.1 |
| HRC Chr10 | 1809068 | 191210 | 1000 | 52330 | 0.1 |
| HRC Chr20 | 829265 | 95414 | 1000 | 52330 | 0.1 |

#### 3.7.1 Simulated Data

We simulated three 10 Mb sequence data sets, with 12,000, 102,000, 1,002,000 haplotypes, using the software `msprime` [16]. We randomly selected 1000 samples (2000 haplotypes) from each pool to form the target genotypes and used the remaining to form the reference panels. All SNPs with minor allele frequency greater than 5% were designated the typed SNPs. Then 0.5% of the typed genotypes were randomly masked to introduce missing values.

#### 3.7.2 1000 Genomes Data

We downloaded the publicly available 1000 Genomes (1000G) phase 3 version 5a data set [1, 2]. This data set contains 2504 samples with 49,143,605 phased genotypes across 26 different populations, as summarized in Table 5 in the supplemental. We focused on chromosomes 10, 18, and 20 data in our experiments. For chromosome 10 and 20, we randomly selected 100 samples to serve as imputation targets and the remaining samples to serve as the reference panel. For chromosome 10 and 20, we chose SNPs also present in the Infinium Omni5-4 Kit to be typed SNPs. We randomly masked 0.1% of the typed genotypes to mimic data missing at random. These data sets feature in our speed and accuracy comparisons. For chromosome 18, we chose the top 50,000 most ancestry informative markers (AIMs) with minor allele frequency ≥ 0.01 as the typed SNPs [5]. The AIM markers were computed using `VCFTools.jl`. To avoid samples with likely substantial degrees of continental-scale admixture, we excluded the populations ACB, ASW, CLM, GIH, ITU, MXL, PEL, PUR, and STU from the reference panel. To illustrate admixture and chromosome painting, our three sample individuals are taken from the excluded populations. The samples from the remaining populations are assumed to exhibit less continental-scale admixture and serve as the reference panel. For admixture analysis, it would be ideal to have samples from the indigenous Amerindian populations as part of the reference panel, but these populations are not surveyed in the 1000 Genomes data, and so we use East Asian (EAS) and South Asian (SAS) populations as the best available proxy.

#### 3.7.3 Haplotype Reference Consortium Data

We also downloaded the Haplotype Reference Consortium (HRC) v1.1 data from the European Genotype-Phenome archive [18] (data accession = EGAD00001002729). This data set consists of 39,741,659 SNPs in 27,165 individuals of predominantly European ancestry. We randomly selected 1000 samples in chromosomes 10 and 20 to serve as imputation targets and the remaining to serve as the reference panel. SNPs also present on the Infinium Omni5-4 Kit are chosen to be typed SNPs. Finally, we randomly masked 0.1% of the typed genotypes to mimic data missing at random.

## 4 Results

Due to Julia’s flexibility, `MendelImpute` runs on Windows, Mac, and Linux operating systems, equipped with either Intel or ARM hardware.

### 4.1 Comparison setup

All programs were run on Linux CentOS 7 equipped with 10 cores of an Intel i9 9920X CPU and 64 GB of RAM. All reference files were previously converted to the corresponding compressed formats, `bref3`, `m3vcf`, `imp5`, or `jlso`. All target genotypes were unphased, and 0.1%-0.5% of typed genotypes were deleted at random. Since `Minimac` 4 and `Impute` 5 required pre-phased data, we used `Beagle` 5.1’s built-in pre-phasing algorithm and report its run-time and RAM usage along side their run times. All output genotypes are phased and complete.

`MendelImpute` was run with Julia v1.5.0. The maximum number of unique haplotypes per window (see supplemental on jlso compression) was set to *d_max_* = 1000, and the number of BLAS threads was set to 1 to avoid over-subscription. Beagle and `Minimac` were run under their default settings. `Impute` 5 was run on 20Mb chunks corresponding to different chromosome regions, except for chromosome 10 of HRC. Chunks were initialized in parallel and imputed separately. Each chunk potentially employs multithreading. We used as many threads as possible without exceeding 10 total threads over all chunks. For chromosome 10 of HRC, we imputed 10Mb chunks, with a maximum of 8 processes active at any given time. Using the maximum 10 processes or employing longer chunks resulted in out of memory error.

### 4.2 Speed, Accuracy, and Peak Memory Demand

Table 3 compares the speed, accuracy, and peak memory (RAM) usage of `MendelImpute`, `Beagle` 5.1, `Impute` 5 version 1.1.4, and `Minimac` 4. Memory was measured via the usr/bin/time command, except for `Impute` 5 where memory is monitored manually via the `htop` command. Note that in our simulated and real data sets the correct values are known for all imputed and masked genotypes. Thus, we can report accuracy as the fraction of genotypes incorrectly imputed for all SNPs, typed or untyped. We do not compute the popular *r*^2^ correlation metric for measuring imputation quality for reasons explained in the supplemental.

**Table 3:** Error, time, and memory comparisons on real and simulated data. The best number in each cell is bolded. Error measurement is possible in these data sets because the correct genotypes are known. `Minimac` 4 and `Impute` 5’s benchmarks do include the pre-phasing step done by `Beagle` 5.1, whose memory and time are reported in brackets. For the sim 1M data, the `m3vcf` reference panel required for `Minimac` could not be computed due to excessive memory requirements.

| **sim 10K** | Error Rate | Time (sec) | Memory (GB) |
| --- | --- | --- | --- |
| MendelImpute | 3.00E-04 | **10** | **1.6** |
| Impute 5 | 2.82E-05 | 43 | 8.9 [6.6] |
| Beagle 5.1 | 2.81E-05 | 189 | 8.8 |
| Minimac 4 | **2.38E-05** | 271 [177] | 1.0 [6.6] |
| **sim 100K** | Error Rate | Time (sec) | Memory (GB) |
| MendelImpute | 2.19E-05 | **14** | **1.6** |
| Impute 5 | 9.36E-06 | 74 [253] | 9.9 [14.5] |
| Beagle 5.1 | 8.22E-06 | 279 | 20.1 |
| Minimac 4 | **7.91E-06** | 3032 [253] | 2.6 [14.5] |
| **sim 1M** | Error Rate | Time (sec) | Memory (GB) |
| MendelImpute | 2.21E-05 | **27** | **4.4** |
| Impute 5 | 1.33E-05 | 153 [752] | 12.6 [25.6] |
| Beagle 5.1 | **7.01E-06** | 769 | 25.6 |
| Minimac 4 | NA | NA | NA |
| **1000G chr20** | Error Rate | Time (sec) | Memory (GB) |
| MendelImpute | 7.63E-03 | **18** | **2.3** |
| Impute 5 | **4.58E-03** | 38 [126] | 7.4 [8.1] |
| Beagle 5.1 | 4.63E-03 | 142 | 8.9 |
| Minimac 4 | **4.58E-03** | 293 [126] | 3.0 [8.1] |
| **1000G chr10** | Error Rate | Time (sec) | Memory (GB) |
| MendelImpute | 7.55E-03 | **35** | **3.3** |
| Impute 5 | **4.83E-03** | 64 [246] | 9.9 [13.2] |
| Beagle 5.1 | 4.91E-03 | 279 | 10.1 |
| Minimac 4 | 4.84E-03 | 588 [246] | 4.2 [13.2] |
| **HRC chr10** | Error Rate | Time (sec) | Memory (GB) |
| MendelImpute | 1.70E-03 | **183** | **8.4** |
| Impute 5 | **8.00E-04** | 754 [2619] | 47.6 [13.0] |
| Beagle 5.1 | 8.34E-04 | 3191 | 33.2 |
| Minimac 4 | 8.57E-04 | 17100 [2619] | 11.5 [13.0] |
| **HRC chr20** | Error Rate | Time (sec) | Memory (GB) |
| MendelImpute | 1.89E-03 | **84** | **5.1** |
| Impute 5 | **8.57E-04** | 645 [1319] | 39.0 [13.3] |
| Beagle 5.1 | 9.10E-04 | 1549 | 22.1 |
| Minimac 4 | 9.18E-04 | 8640 [1319] | 15.1 [13.3] |

On HRC, `MendelImpute` runs 18-23 times faster than `Impute` 5 (including pre-phasing time), 17-18 times faster than `Beagle` 5.1, and 108-119 times faster than `Minimac` 4. On the smaller 1000 Genomes data set, `MendelImpute` runs 9 times faster than `Impute` 5, 8 times faster than `Beagle` 5.1, and 23-24 times faster than `Minimac` 4. Increasing the reference panel size by a factor of 100 on simulated data only increases `MendelImpute`’s computation time by a factor of at most three. `MendelImpute` also scales better than HMM methods as the number of typed SNPs increases. Thus, denser SNP arrays may benefit disproportionately from using `MendelImpute`. The 1000 Genomes data set is exceptional in that it has fewer than 5000 reference haplotypes. Therefore, traversing that HMM state space is not much slower than performing the corresponding linear algebra calculations in `MendelImpute`. Notably, except for the HRC panels, `MendelImpute` spends at least 50% of its total compute time importing data.

In terms of error rate, `MendelImpute` is 1.5-2.8 times worse on simulated and real data than `Impute` 5, `Minimac` 4, and `Beagle` 5.1. The sim10k data set is an exception in that `MendelImpute`’s error rate is 10 times worse, which we attribute to the size of the reference panel compared to the number of imputed samples. The error rates of `Impute` 5, `Beagle` 5.1, and `Minimac` 4 are similar, consistent with previous findings [24]. As discussed in the supplemental, it is possible to improve `MendelImpute`’s error rate by more computationally intensive strategies such as phasing by dynamic programming.

Finally, `MendelImpute` requires much less memory for most data sets. As explained in the methods section and the supplemental, the genotype matrix and compressed reference panel are compactly represented in memory. Since most analysis is conducted in individual windows, only small sections of these matrices need to be decompressed into single-precision arrays at any one time. Consequently, `MendelImpute` uses at most 8.4 GB of RAM in each of these experiments. In general, `MendelImpute` permits standard laptops to conduct imputation even with the sizeable HRC panels.

### 4.3 Local Ancestry Inference for Admixed Populations

`MendelImpute` lends itself to chromosome painting, the process of coloring each haplotype segment by the country, region, or ethnicity of the reference individual assigned to the segment. For chromosome painting to be of the most value, reference samples should be from individuals who are representative of distinct populations. Within a reference population there should be little admixture. Also the colors assigned to different regions should be coordinated by physical, historical, and ethnic proximity. The overall proportions of the colors assigned to a sample individual genome immediately translate into admixture coefficients. Here we illustrate chromosome painting using chromosome 18 data from the 1000 Genomes Project. The much larger Haplotype Reference Consortium data would be better suited for chromosome painting, but unfortunately its repository does not list country of origin. Our examples should therefore be considered simply as a proof of principle. As already mentioned, the populations present in the 1000 Genomes Project data are summarized in Table 5 of the supplemental.

Figure 2A displays the painted chromosomes 18 of a native Puerto Rican (PUR, sample 1), a Peruvian from Lima, Peru (PEL, sample 2), and a person of African ancestry from the Southwest USA (ASW, sample 3). Here a total of 17 reference populations potentially contribute genetic segments. They are colored with red, brown, blue, or green to capture South Asian, East Asian, European, or African backgrounds, respectively. Note that the samples from South/East Asian populations serve as a proxy for Amerindian ancestral populations. After coloring, the two PUR extended haplotypes are predominantly blue, the two PEL haplotypes are predominantly red/brown and blue, while the two ASW haplotypes are predominantly green. Interestingly, one of the PUR and one of the PEL haplotypes contain a block of African origin as well as blocks of Asian and European origin, while the ASW haplotypes contain two blocks of European origin. The relatively long blocks are suggestive of recent admixture. The resulting chromosome barcodes vividly display population origins and suggest the locations of ancient or contemporary recombination events.

**Figure 2:**
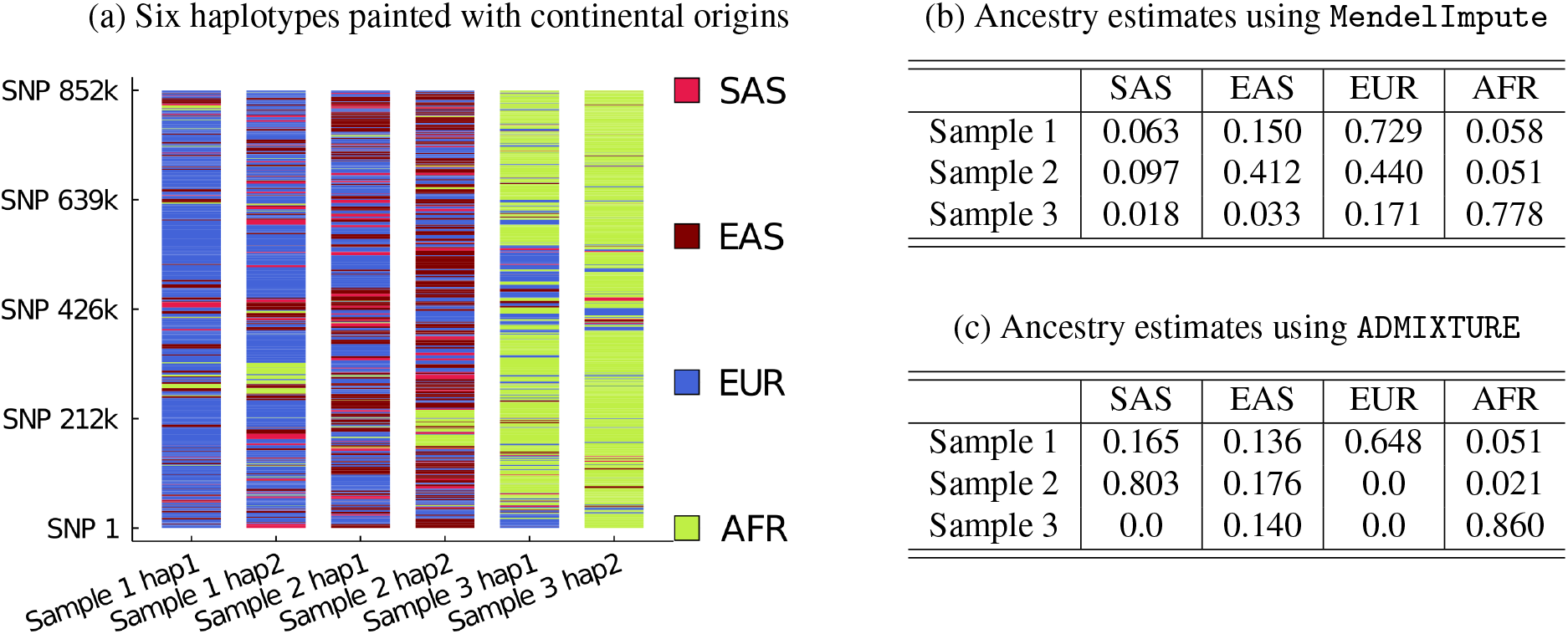
`MendelImpute` used for chromosome painting and ancestry estimation. (a) Painted chromosome 18 after phasing for three samples. Sample 1 is Puerto Rican (PUR), sample 2 is Peruvian (PEL), and sample 3 is African American (ASW). In the reference populations, South Asians are shaded in red, East Asians in brown, Europeans in blue, and Africans in green. (b) Global ancestry proportions estimated from the cumulative length of the colored segments in (a). (c) Global ancestry estimates using the `ADMIXTURE` software.

Figure 2B displays the three samples’ admixture proportions estimated from the cumulative lengths of each color in Figure 2A. We compared `MendelImpute`’s estimates to our ADMIXTURE software [3], whose results are listed in Figure 2C. Samples 1 and 3 match well with `MendelImpute`. For sample 2, ADMIXTURE estimates roughly 98% of South/East Asian ancestry, disagreeing with `MendelImpute` which estimates roughly 50-50 split between South/East Asian and European ancestry. Of course, in actual practice we would use all chromosomes of an individual, which would provide a far more accurate assessment than using just chromosome 18. In addition, the ancestral assignments are only as good as the choice of reference haplotypes. For instance, the haplotypes of the African American (sample 3; ASW) contain a small but substantial portion (3%) of Eastern Asian ancestry. This labelling should not be taken too literally but rather may reflect either some distant Amerindian ancestry or regions where many haplotypes are similar and so ancestry is difficult to discern.

### 4.4 Ultra-Compressed Phased Genotype Files

As discussed earlier, VCF files are enormous and slow to read. If genotypes are phased with respect to a particular reference panel, then an alternative is to store each haplotype segment’s starting position and a pointer to its corresponding reference haplotype. This offers massive compression because long haplotype segments are reduced to two integers. Instead of the default compressed VCF files, `MendelImpute` can optionally output such ultra-compressed phased data. Table 4 shows that the ultra-compressed format gives 20-270 fold compression compared to standard compressed VCF outputs. In principle, all phased genotypes can be stored in such files. The drawback is that compressed data can only be decompressed with the help of the original reference panel. Thus, this tactic relies on universal storage and curation of reference haplotype panels. These panels should be stored on the cloud for easy access and constructed so that they can be consistently augmented by new reference haplotypes.

**Table 4:**
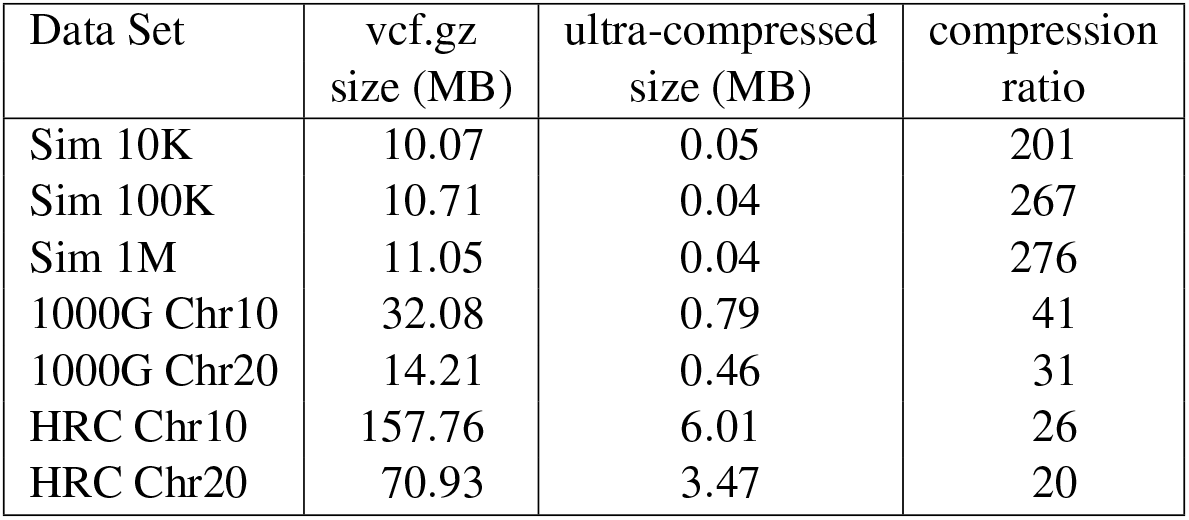
Output file size comparison of compressed VCF and ultra-compressed formats.

| Data Set | vcf.gz<br>size (MB) | ultra-compressed<br>size (MB) | compression<br>ratio |
| --- | --- | --- | --- |
| Sim 10K | 10.07 | 0.05 | 201 |
| Sim 100K | 10.71 | 0.04 | 267 |
| Sim 1M | 11.05 | 0.04 | 276 |
| 1000G Chr10 | 32.08 | 0.79 | 41 |
| 1000G Chr20 | 14.21 | 0.46 | 31 |
| HRC Chr10 | 157.76 | 6.01 | 26 |
| HRC Chr20 | 70.93 | 3.47 | 20 |

**Table 5:** The 26 population codes present in the 1000 genomes project.

| Population Code | Description | Super Population |
| --- | --- | --- |
| CHB | Han Chinese in Beijing, China | East Asian (EAS) |
| JPT | Japanese in Tokyo, Japan | East Asian (EAS) |
| CHS | Southern Han Chinese | East Asian (EAS) |
| CDX | Chinese Dai in Xishuangbanna, China | East Asian (EAS) |
| KHV | Kinh in Ho Chi Minh City, Vietnam | East Asian (EAS) |
| CEU | Utah Residents with NW European Ancestry | European (EUR) |
| TSI | Toscani in Italia | European (EUR) |
| FIN | Finnish in Finland | European (EUR) |
| GBR | British in England and Scotland | European (EUR) |
| IBS | Iberian Population in Spain | European (EUR) |
| YRI | Yoruba in Ibadan, Nigeria | Africans (AFR) |
| LWK | Luhya in Webuye, Kenya | Africans (AFR) |
| GWD | Gambian in Western Divisions in the Gambia | Africans (AFR) |
| MSL | Mende in Sierra Leone | Africans (AFR) |
| ESN | Esan in Nigeria | Africans (AFR) |
| ASW | Americans of African Ancestry in SW USA | Africans (AFR) |
| ACB | African Caribbeans in Barbados | Africans (AFR) |
| MXL | Mexican Ancestry from Los Angeles USA | Ad Mixed American (AMR) |
| PUR | Puerto Ricans from Puerto Rico | Ad Mixed American (AMR) |
| CLM | Colombians from Medellin, Colombia | Ad Mixed American (AMR) |
| PEL | Peruvians from Lima, Peru | Ad Mixed American (AMR) |
| GIH | Gujarati Indian from Houston, Texas | South Asian (SAS) |
| PJL | Punjabi from Lahore, Pakistan | South Asian (SAS) |
| BEB | Bengali from Bangladesh | South Asian (SAS) |
| STU | Sri Lankan Tamil from the UK | South Asian (SAS) |
| ITU | Indian Telugu from the UK | South Asian (SAS) |

## 5 Discussion

We present `MendelImpute`, the first scalable, data-driven method for phasing and genotype imputation. `MendelImpute` and supporting `OpenMendel` software [28] provide an end-to-end analysis pipeline in the Julia programming language that is typically 10–100 times faster than methods based on hidden Markov models, including `Impute` 5, `Beagle` 5.1 (Java) and `Minimac` 4 (C++). The speed difference increases dramatically as we increase the number of typed SNPs. Thus, denser SNP chips potentially benefit more from `MendelImpute`’s design. Furthermore, `MendelImpute` occupies a smaller memory footprint. This makes it possible for users to run `MendelImpute` on standard desktop or laptop computers on very large data sets such as the HRC. Unfortunately we cannot yet have the best of both worlds, as `MendelImpute` exhibits a 1.5-2.2 fold worse error rate on real data. However, as seen in Table 3, `MendelImpute`’s error rate is still acceptably low. One can improve its error rate by implementing strategies such as phasing by dynamic programming. This strategy may decrease error, but it slows down computation and is hence omitted in our comparisons. Regardless, it is clear that big data methods can compete with HMM based methods on the largest data sets currently available and that there is still room for improvement and innovation in genotype imputation.

Beyond imputation and phasing, our methods extend naturally to ancestry estimation and data compression. If each reference haplotype is labeled with its country or region of origin, then `MendelImpute` can decompose a sample’s genotypes into segments of different reference haplotypes colored by these origins. The cumulative lengths of these colored segments immediately yield an estimate of admixture proportions. Countries can be aggregated into regions if too few reference haplotypes originate from a given country. The colored segments also present a chromosome barcode that helps one visualize subject variation, recombination hotspots, and global patterns of linkage disequilibrium. Data compression is achieved by storing the starting positions of each segment and its underlying reference haplotype. This leads to output files that are 20-270 fold smaller than standard compressed VCF files. Decompression obviously requires ready access to stable reference panels stored on accessible sites such as the cloud. Although such an ideal resource is currently part dream and part reality, it could be achieved by a concerted international effort.

For potential users and developers, the primary disadvantage of `MendelImpute` is its reliance on the importation and storage of a haplotype reference panel. Acquiring these panels requires an application process which can take time to complete. Understanding, storing, and wrangling a panel add to the burden. The imputation server for `Minimac` 4 thrives because it relieves users of these burdens [11]. `Beagle` 5.1 is capable of fast parallel data import on raw VCF files [7], which neither `Minimac` 4 nor `MendelImpute` can currently match. This makes target data import, and especially pre-processing the reference panel, painfully slow for both programs. Fortunately, pre-processing only has to happen once.

Finally, let us reiterate the goals and achievements of this paper. First, we show that data-driven methods are competitive with HMM methods on genotype phasing and imputation, even on the largest data sets available today. Second, we challenge the notion that pre-phasing and imputation should be kept separate; `MendelImpute` performs both simultaneously. Third, we argue that data-driven methods are ultimately more flexible; for instance, `MendelImpute` readily handles imputation and phasing on dosage data. Fourth, we demonstrate that data-driven methods yield dividends in ancestry identification and data compression. Fifth, `MendelImpute` is completely open source, freely downloadable, and implemented in Julia, an operating system agnostic, high-level programming language for scientific research. Julia is extremely fast and enables clear modular coding. Our experience suggests that data-driven methods will offer a better way forward as we face increasingly larger reference panels, denser SNP array chips, and greater data variability.

## 6 Supplemental

### 6.1 Imputation Quality Scores

Consider the observed genotype *x_ij_* ∈ [0, 2] ∪ {*missing*} at SNP *i* of sample *j* and the corresponding imputed genotype *g_ij_* derived from the two extended haplotypes of *j*. If *S_i_* denotes the set of individuals with observed genotypes at the SNP, then `MendelImpute`’s quality score *q_i_* for the SNP is defined as

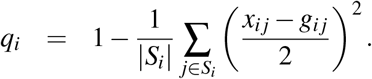

Note that 0 ≤ *q_i_* ≤ 1 and that the larger the quality score, the more confidence in the imputed values. Because *q_i_* can only be computed for the typed SNPs, an untyped SNP is assigned the average of the quality scores for its two closest flanking typed SNPs. Figure 3A plots each SNP’s quality score in the 1000G Chr20 experiment summarized in Table 3. For each sample, one can also compute the mean least squares error over all *p* SNPs to obtain a per-sample quality score. This is shown in Figure 3B. By default `MendelImpute` outputs both quality scores. Thus, investigators can perform post-imputation quality control by SNPs and by samples separately.

**Figure 3:**
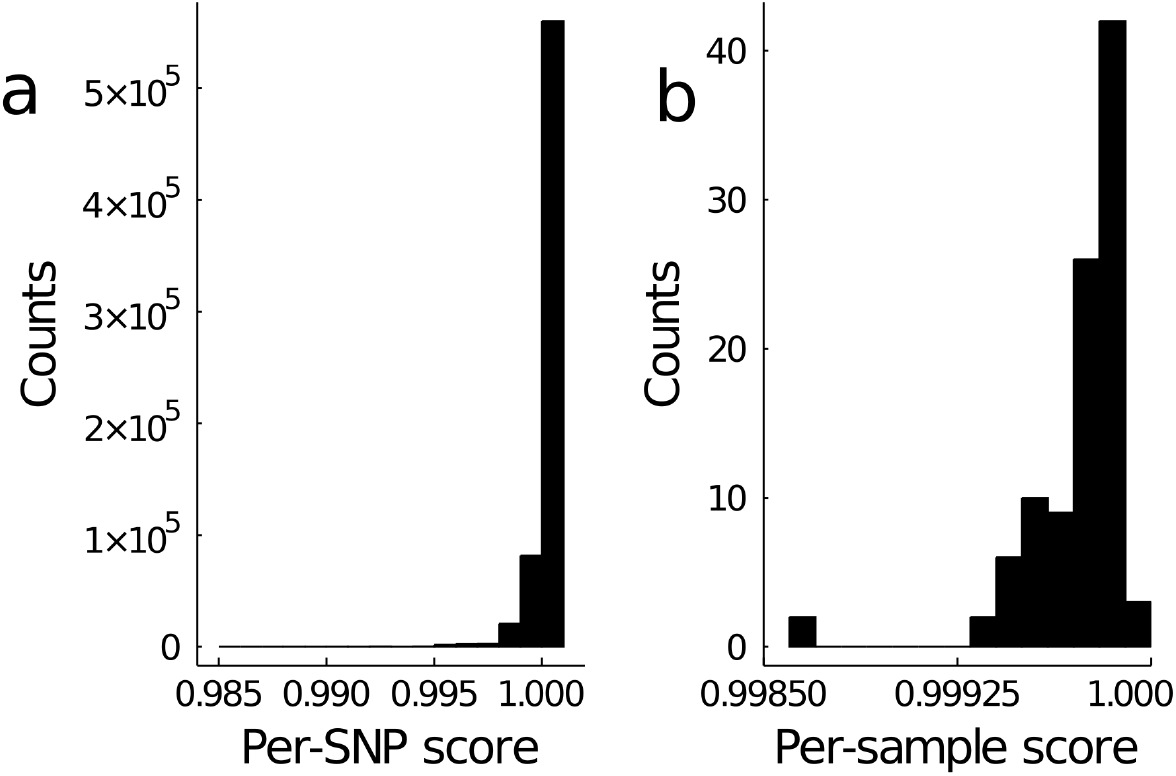
Histograms of per-SNP and per-sample quality scores for chromosome 20 in our 1000G analysis. By default `MendelImpute` computes (a) per-SNP quality scores and (b) per-sample quality scores. SNPs and samples with noticeably lower quality scores should be removed from downstream analysis.

Empirically, it is rather common for a sample subject to harbor a few poorly imputed windows. Thus, we observe a long left tail in the histogram for per-sample error in Figure 3. Unfortunately, the bad windows generally do not exhibit any discernible regional patterns across subjects. We suspect that poorly imputed windows involve breakpoints that occur near the middle of a window. We plan a detailed analysis of this issue in future work.

#### 6.1.1 Imputation quality scores using r-squared

As mentioned previously, the popular *r*^2^ correlation coefficient between imputed genotypes and true genotypes is an uninformative metric for measuring imputation quality for `MendelImpute`. Although true genotypes are unknown, *r*^2^ can still be estimated from posterior probabilities of the HMM model [6]. For instance, according to `Minimac` 3’s online documentation, for each SNP we can calculate

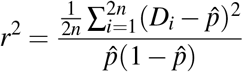

where 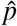 is the alternative allele frequency (of the imputed data), *D_i_* is the imputed alternate allele probability at the *i*th haplotype, and *n* is the number of GWAS samples. `MendelImpute` does not compute posterior probabilities; for each locus, each haplotype is imputed with 0 or 1. That is, *D_i_* ∈ {0, 1}. Thus, using the fact that 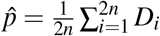 we have

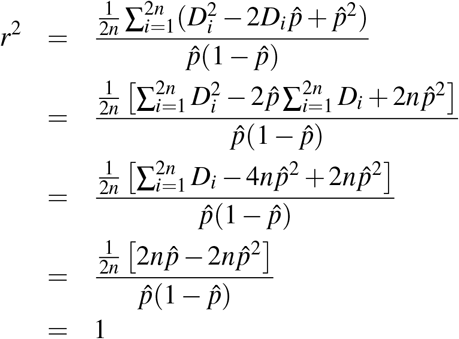

Thus every SNP has *r*^2^ = 1 under `MendelImpute`’s model, which renders it meaningless.

### 6.2 JLSO Compressed Reference Haplotype Panels

The `jlso` format is constructed in three steps: (a) specify window intervals, (b) compute unique haplotypes in addition to hash maps to reference haplotypes in each window, and (c) save the result in a binary compressed format via the `JLSO.jl` package [12]. The resulting `jlso` files are 30-50x faster to read and 3-5x smaller in file size (varies depending on window width) than compressed VCF files in the `vcf.gz` format. Note the `jlso` format is simply a container object that facilitates reading and transferring large VCF files stored as Julia variables. In principle, all files that are slow to read can be pre-processed and stored in this alternative format for quicker access. As such, we similarly store ultra-compressed sample haplotypes, as discussed in Section 4.4, using the `JLSO.jl` package.

#### 6.2.1 Adaptive Window Widths via Recursive Bisection

The first step in generating `JLSO` haplotype panel is to specify genomic window ranges. The width of genomic windows is an important parameter determining both imputation efficiency and accuracy. Empirically, larger window widths give better error rates but also increase the computational burden of the matrix multiplications and minimum entry search described in Section 3.2. The magnitudes of these burdens depend on local haplotype diversity. Thus, we choose window widths dynamically. This goal is achieved by a bisection strategy. After aligning all typed SNPs with the reference panel, initially we view all typed SNPs on a large section of a chromosome as belonging to a single window. We then divide the window into equal halves if it possesses too many unique haplotypes. Each half is further bisected and so forth recursively until every window contains fewer than a predetermined number of unique haplotypes. Empirically, choosing the maximum number *d_max_* of unique haplotypes per window to be 1000 works well for both real and simulated data. When a larger number is preferred, we resort to a stepwise search heuristic for minimizing criterion (3.1) that scales linearly in the number of unique haplotypes *d*. This heuristic is described above.

#### 6.2.2 Elimination of Redundant Haplotypes by Hashing

Within a small genomic window of the reference panel, multiple haplotype pairs may be identical at the typed SNPs. Only the unique haplotypes play a role in matching reference haplotypes to sample genotypes. `MendelImpute` identifies redundant haplotypes by hashing. For each reference haplotype limited to the window, hashing stores an integer representation of the haplotype via a hash function. This integer serves as an index (key) to locate the reference haplotype (value). Put another way, hashing stores the inverse images of the map from reference haplotypes to unique haplotypes. In our software, the `GroupSlices.jl` package [13] identifies a unique key for each haplotype.

#### 6.2.3 Save in binary compressed format

Since haplotypes are long binary vectors, the entire haplotype reference panel can be compactly represented using a single bit per entry. We have already divided the full reference panel into non-overlapping windows of various widths after proper alignment of all typed and reference SNPs in the window. For each window we save two compressed mini-panels. The first houses the unique haplotypes determined by just the typed SNPs. The second houses the unique haplotypes determined by all SNPs in the window, typed or untyped. The former is much smaller than the later, but each entry of both can be compactly represented by a single bit per entry in memory. Thus, for each window we save two compressed windows in addition to meta information and pointers that coordinate reference haplotypes with the two mini-panels per each window. This whole ensemble is stored in the jlso file. Because there are only a limited number of SNP array chips on the market, one can in principle store just a few jlso files on a universal source such as the cloud.

### 6.3 Parallel Computing and Memory Requirements

`MendelImpute` employs a shared-memory parallel computing model where each available core handles an independent component of the entire problem. Work is assigned via Julia’s multi-threading functionality. When computing the optimal haplotype pairs in equation (3.1), we parallelize over windows. This requires allocating *c* copies of **X**^*T*^ **H** and **H**^*T*^ **H**, where *c* is the number of CPU cores available. Note the dimensions of these matrices vary across windows. To avoid accruing memory allocations, we pre-allocate *c* copies of *n* × *d_max_* and *d_max_* × *d_max_* matrices and re-use their top-left corners in windows with *d < d_max_*. For intersecting adjacent reference haplotype sets (phasing), we parallelize over samples. This step requires no additional memory. Writing to output is also trivially parallelizable by assigning each thread to write a different portion of the imputed matrix to a different file, then concatenating these files into a single output file. Data import is not parallelized. Beyond allocating **X**^*T*^ **H** and **H**^*T*^ **H**, our software requires enough memory (RAM) to load the target genotype matrix and the compressed haplotype reference panel.

### 6.4 Bias Correction for Initializing Missing Data

Since BLAS requires complete data, we must first initialize the missing data in each genotype vector **x** before computing **M** and **N** in equation (3.2). This may introduce bias in our minimization of criterion (3.1) if there is a high fraction of missing genotypes in the typed SNPs, for example above 10%. One way to alleviate bias is to initialize missing data with the mean and save all unique haplotype pairs minimizing criterion (3.1) under this convention. Once this set of optimal haplotype pairs are identified, we re-minimize criterion (3.1) but now skipping the missing entries of **x**. That is equivalent to setting *x_k_* − *h_ik_* − *h _jk_* = 0 when *x_k_* is missing.

### 6.5 Avoidance of Global Searches for Optimal Haplotype Pairs

Recall that minimizing the criterion (3.1) requires searching through all lower-triangular entries of the *d* × *d* matrix **M** + **N**, where *d* denotes the number of unique haplotypes in the window. When *d <* 1000, searching through all 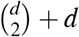 lower-triangular entries of **M** + **N** via `MendelImpute`’s standard procedure is fast, but this global search quickly degrades as *d* → ∞. Below we outline two heuristic procedures for large *d*. These heuristics typically produce sub-optimal solutions compared to global searches, so they should be used with caution.

#### 6.5.1 Stepwise Search Heuristics

Consider minimizing the loss 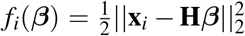, where the *d* columns of **H** ∈ {0, 1}^*p*×*d*^ store unique haplotypes, *p* is the window width, and **x**_*i*_ is a sample genotype vector. The original problem (3.1) minimizes *f_i_*(****β****) under the constraint that exactly two *β_j_* = 1 and the remaining *β_k_* = 0 or the constraint that exactly one *β_j_* = 2 and the remaining *β_k_* = 0. As an approximate alternative, one first finds the *r* unique haplotypes with the largest influence o n *f _i_*(****β****). This i s accomplished by identifying the *r* most negative components of the gradient

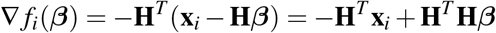

at ****β**** = **0**. These are the *r* directions of steepest descent. Note that ∇ *f_i_*(**0**) = −**H**^*T*^ **x**_*i*_ and that **H**^*T*^ **x**_*i*_ is pre-computed and cached in **N**. The residual function 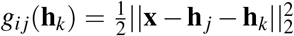 is then minimized over **h**_*k*_ to find the candidate pair (**h**_*j*_, **h**_*k*_) generated by each of the vectors **h**_*j*_ determined by the gradient ∇ *f_i_*(**0**). The ingredients to perform these minimizations are already in hand. This heuristic scales as 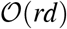, much better than 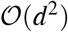 in equation (3.2). `MendelImpute` sets the default *r* = 100. In the same spirit as the first step, one can alternatively find for each *j* the most negative component *k* of the gradient ∇ *f_i_*(**e**_*j*_), where **e**_*j*_ is the standard unit vector with 1 in position *j*. This again determines a nearly optimal pair (**h**_*j*_, **h**_*k*_). Under this tactic the Gram matrix **H**^*T*^ **H** comes into play. Note that **H**^*T*^ **He**_*j*_ reduces to its *j*th column **v**_*j*_. Hence, no new matrix-by-vector multiplications are necessary in calculating ∇ *f_i_*(**e**_*j*_) = −**H**^*T*^ **x**_*i*_ + **v**_*j*_ = ∇ *f_i_*(**0**) + **v**_*j*_.

Alternatively, one can find the best *r* unique haplotypes for a given sample **x**_*i*_ en masse by arranging all pairwise column distances of **X** and **H** in the matrix

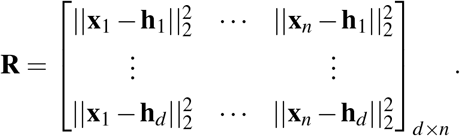

Then we partially sort each column of **R** to identify the top *r* haplotypes matching each sample **x**_*i*_. Here **R** is computed via the Distance.jl package of Julia, which internally performs BLAS level-3 calls analogous to computing **H**^*T*^ **H** and **X**_*T*_ **H**. Instead of searching through all haplotypes to minimize *g_i__j_*(**h**_*k*_) for a given sample **x**_*i*_, one can instead search only over the 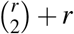 combinations of the top haplotypes. This allows one to entertain much larger values of *r*. Empirically, choosing *r* = 800 works well for most data sets.

### 6.6 Phasing by Dynamic Programming

We also investigated a dynamic programming strategy that gives the global solution for minimizing the number of haplotype breaks across the extended haplotypes **E**_1_ and **E**_2_. For each given haplotype pair **p**_1_ = (**h**_*i*_, **h**_*j*_) in window *w*, we can compute the squared Hamming distance between it and the pair **p**_2_ = (**h**_*k*_, **h**_*l*_) in window *w* + 1; in symbols

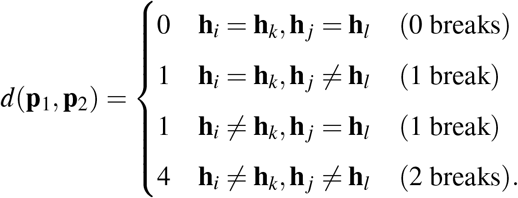

Observe that a double break is assigned an error of 4 to favor 2 single breaks across 3 windows as opposed to a double break plus a perfect match.

Now we describe a dynamic programming strategy for finding the two paths with the minimal number of unique haplotype breaks. We start with all candidate pairs **p**_*i*_ in the leftmost window and initialize sums *s_i_* = 0 and traceback path vectors **t**_*i*_ to be empty. One then recursively visits all windows in turn from left to right. If *w* is the current window, then every candidate haplotype pair **p**_*i*_ in window *w* is connected to every candidate pair **p**_*j*_ in window *w* + 1. The traceback path **t**_*j*_ is determined by the pair **p**_*k*_ minimizing *d*(**p**_*i*_, **p**_*j*_). The traceback path **t**_*j*_ is constructed by appending **p**_*k*_ to **t**_*k*_ and setting *s _j_* = *s_k_* + *d*(**p**_*k*_, **p**_*j*_). This process is continued until the rightmost window *v* is reached. At this point the pair **p**_*j*_ with lowest running sum *s _j_* is declared the winner. The traceback path **t**_*j*_ allows one to construct the extended haplotypes **E**_1_ and **E**_2_ in their entirety. Unfortunately, too many haplotype pairs per window can overwhelm dynamic programming with large reference panels. One partial recourse is to discard partial paths and associated partial termini **p**_*i*_ that are unpromising in the sense that their running sums *s_i_* are excessively large. This is effective, but the approximate algorithm is burdened by extra bookkeeping. The bookkeeping of the exact algorithm is already demanding.

### 6.7 Summary of 1000 Genomes Reference Panel

A total of 26 different populations contribute to the 1000 Genomes Project data set. These populations are further organized into five super population. While this information is freely available online, we summarize it in Table 5 for completeness.

## 7 Author contributions

KL, JS, ES, and HZ conceived this project. BC, KL, RW, JS, ES, and HZ devised the methods. BC, RW, and HZ developed the software. BC and KL accessed the data. BC wrote the original draft of the paper. BC, KL, RW, JS, ES, and HZ reviewed and edited the draft. KL previewed some of the methods incorporated in `MendelImpute` in his talk [17].

## 8 Acknowledgements

BC and RW were supported by NIH grant T32-HG002536. BC, ES, JS, HZ, and KL were supported by NIH grant R01-HG006139. ES, JS, HZ, and KL were supported by NIH grant R01-GM053275. JS was also supported by NIH grant R01-HG009120.

We would also like to thank Calvin Chi for his helpful discussions on ancestry estimation and Juhyun Kim for her helpful discussions on imputation quality scores.

## 10 Web Resources

**Project name**: MendelImpute.jl

**Project home page**: https://github.com/OpenMendel/MendelImpute.jl

**Supported operating systems**: Mac OS, Linux, Windows

**Programming language**: Julia 1.5

**License**: MIT

All commands needed to reproduce the following results are available at the `MendelImpute` site in the manuscript sub-folder. `SnpArrays.jl` is available at https://github.com/OpenMendel/SnpArrays.jl. `VCFTools.jl` is available at https://github.com/OpenMendel/VCFTools.jl. The Haplotype Reference Consortium data is available at https://www.ebi.ac.uk/ega/datasets/EGAD00001002729. The 1000 genomes data is available at ftp://ftp.1000genomes.ebi.ac.uk/vol1/ftp/release/20130502/.

